## Supplementary material for "Highly expressed miR-375 is not an intracellular oncogene in Merkel cell polyomavirus-associated Merkel cell carcinoma": Suppl. Fig

### Suppl. Figure S1

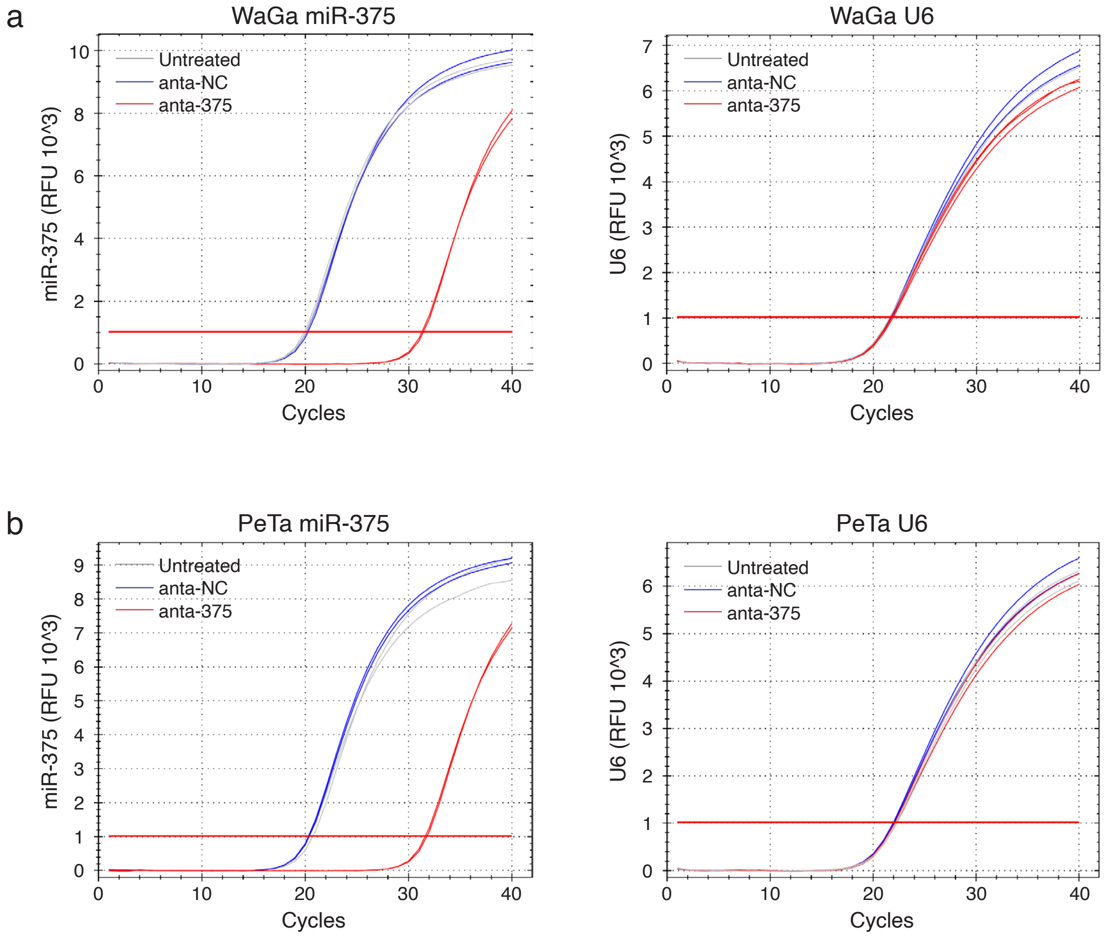


**Supplementary figure S1**: Expression of miR-375 in WaGa (a) and PeTa cells (b) transfected with miR-375 antagomiRs (anta-375, red) or negative control (anta-NC, blue) as well as untransfected cells (Untreated, gray) was determined by RT-qPCR, U6 is used as endogenous control. Depicted are the amplification curves of miR-375 and U6 in duplicates.

### Suppl. Figure S2

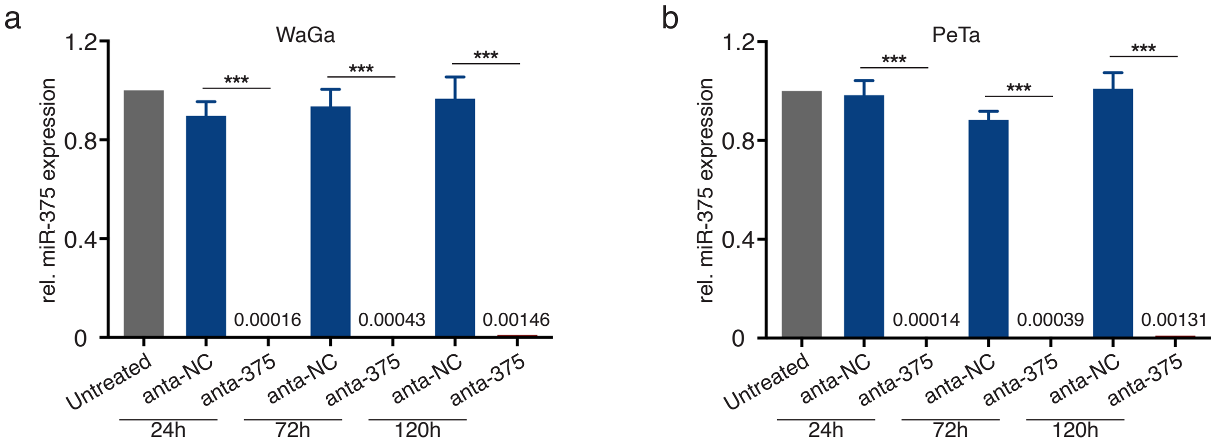


**Supplementary figure S2**: Relative miR-375 expression was determined in triplicates by RT-qPCR in WaGa (a) and PeTa (b) cells transfected with miR-375 antagomiRs (anta-375, red) or negative control (anta-NC, blue) as well untransfected cells (Untreated, gray). Ct values were normalized to U6 and calibrated to the untreated cells. Numbers depicted are relative miR-375 expression values in cells transfected with miR-375 antagomiRs comparing to untreated cells. All experiments were independently repeated three times. Error bars represent SD, *** indicates p<0.001..

### Suppl. Figure S3

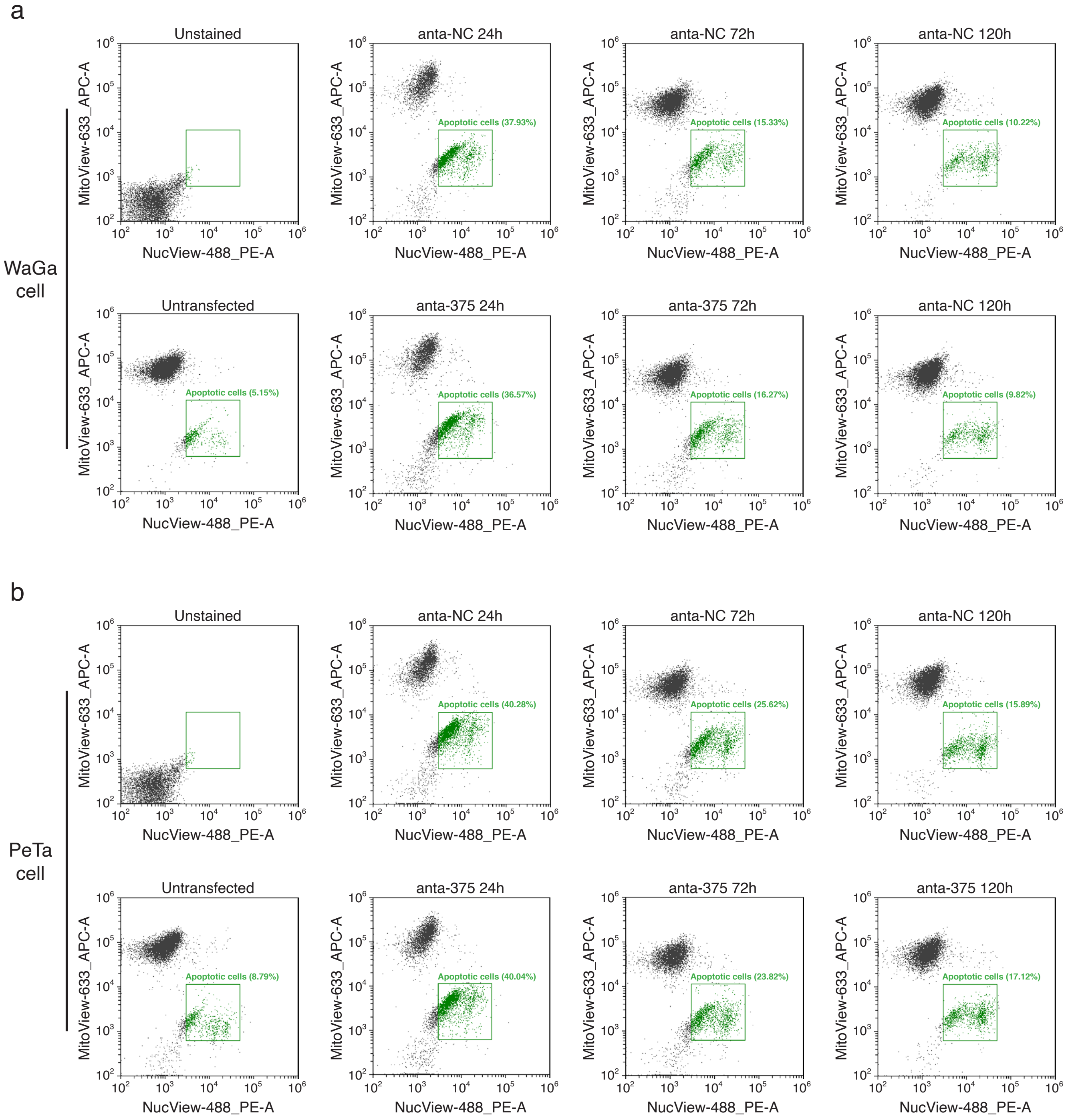


**Supplementary figure S3**: NucView 488/MitoView 633 Apoptosis Assay Kit was applied to determine cell apoptosis of WaGa (a) and PeTa (b) cells transfected with miR-375 antagomiRs (anta-375) or negative control (anta-NC) in flow cytometry. Healthy cells are stained with MitoView 633 (red, APC-A channel), while late apoptotic cells are stained with NucView 488 (green, PE-A channel). Representative flow cytometry dot plots are depicted to reveal apoptotic cell rates.
